# Common anti-haemostatic medications increase the severity of systemic infection by uropathogenic *Escherichia coli*

**DOI:** 10.1101/2021.01.18.425391

**Authors:** Vi LT Tran, Elinor Hortle, Warwick J Britton, Stefan H Oehlers

## Abstract

Uropathogenic *Escherichia coli* (UPEC) causes urinary tract infections that can result in sepsis. The haemostatic system is protective in the pyelonephritis stage of ascending UPEC infection, but the role of the haemostatic system has not been investigated during sepsis. Here we utilize a zebrafish-UPEC systemic infection model to visualize infection-induced coagulation and examine the effects of commonly prescribed anti-haemostatic medications on the infection severity. Treatment of systemically infected zebrafish with warfarin, aspirin, or ticagrelor reduced host survival, while stabilization of clots with aminocaproic acid increased host survival. Anti-haemostatic drug treatment increased UPEC burden. Our findings provide evidence that commonly prescribed anti-haemostatic medications may worsen the outcome of severe UPEC infection.

## Introduction

Uropathogenic *Escherichia coli* (UPEC) is the primary pathogen responsible for urinary tract infection (UTI), which is among the most common bacterial infections worldwide. UPEC most commonly initiates self-limiting cystitis inside the urinary tract, but can ascend the ureters to the kidneys as acute pyelonephritis and potentially progress into sepsis (Wiles, Kulesus et al. 2008, Ulett, Totsika et al. 2013).

The haemostatic system plays an evolutionarily conserved role in both immune control of infections and the pathogenesis of infectious complications from a wide range of pathogens (Frick, Bjorck et al. 2007, Stroo, Zeerleder et al. 2017, Hortle, Johnson et al. 2019). In the case of UPEC, treatment of rodent UPEC pyelonephritis models with the injectable anticoagulant heparin demonstrates an important role of coagulation in preventing the progression of pyelonephritis to sepsis via an α-hemolysin/renal epithelial cell CD147/Tissue Factor axis (Melican, Boekel et al. 2008, Schulz, Chuquimia et al. 2018).

The prevalence of urinary tract infections rises from ∼10% in the general population of women to ∼20% in women over 65 (Medina and Castillo-Pino 2019). Catheterization further increases the risk of urinary tract infections, with a 10-25% risk of a catheter-associated urinary tract infection (Medina and Castillo-Pino 2019). The rise in prevalence of chronic cardiovascular conditions, such as atrial fibrillation (Alcusky, McManus et al. 2019), causes a strong correlation between age and the use of anti-haemostatic medications which might compound the risk and severity of UPEC urosepsis in the elderly. Here, we have used a larval zebrafish model of systemic UPEC infection to study the interaction between commonly prescribed anti-haemostatic medications and the severity of UPEC sepsis (Wiles, Bower et al. 2009).

## Materials and methods

### Zebrafish husbandry

Adult zebrafish were housed at the Centenary Institute (Sydney Local Health District AWC Approval 2017-036). Zebrafish embryos were produced by natural spawning and raised at 28°C in E3 media.

### Zebrafish lines

Wild type zebrafish are the AB background. Transgenic lines are: *Tg(fabp10a:fgb-EGFP)*^*mi4001*^ which was used to visualize clot formation (Vo, Swaroop et al. 2013), and *Tg(−6*.*0itga2b:eGFP)*^*la2*^ which was used to visualize thrombocytes (Lin, Traver et al. 2005).

### Infection of zebrafish larvae

Stationary phase *Escherichia coli* MG-1655 or UTI89 carrying the pGI6 plasmid were outgrown in LB broth supplemented with 50 µg/ml spectinomycin for three hours at 37°C. Bacteria were pelleted, washed in PBS, and either resuspended in phenol red dye (0.5% w/v in PBS) for immediate microinjection or supplemented with 10% v/v glycerol and frozen at −80°C (Wright, de Silva et al. 2021). 10-15 nL was injected into the caudal vein or trunk of M-222 (tricaine)-anaesthetized 5 dpf larvae resulting in infectious doses as reported. Larvae were recovered into E3 and housed at 32°C.

### CFU recovery assay

Groups of 5 zebrafish larvae were pooled and homogenized by pipetting through a P200 tip followed by shearing through 23 G and 28 G needles. Homogenate was serially diluted and plated on LB agar supplemented with 50 µg/ml spectinomycin to select *E. coli* carrying the pGI6 plasmid.

### Drug treatments

Larvae were treated by immersion exposure to vehicle control (DMSO or water as appropriate), 10 µg/ml aspirin in 0.1% final DMSO, 20 µg/ml ticagrelor in 0.1% final DMSO, 20 µM warfarin in water, or 100 mM amino caproic acid (ACA) in water immediately after infection. Drugs were refreshed after two days only for the survival experiments lasting longer than two days, all other experiments were performed with a single dose.

### Imaging

Imaging was carried out on larvae anaesthetized in M-222 mounted in 0.75% low melting point agarose on a Deltavision Elite fluorescence microscope for 24 hours. Editing and bacterial fluorescent pixel count was carried out with Image J Software Version 1.51j (Matty, Oehlers et al. 2016).

### Wound haemostasis assay

We transected the tails of M-222 anesthetized larvae with a scalpel at the ventral pigment gap. This severs the dorsal aorta and posterior cardinal vein resulting in rapid haemostasis in control larvae. Larvae were recovered to E3 prior to imaging at 2 hours post wounding.

### Statistics

Survival analyses were performed by Log-rank tests in GraphPad Prism. Fluorescent pixel count analyses were performed by Student’s *t*-test or ANOVA in GraphPad Prism as appropriate. Error bars represent standard error of the mean.

## Results

### Characterization of infection kinetics in zebrafish larvae

Prior reports extra intestinal *E. coli* infection have largely used embryo stage zebrafish prior to 3 dpf, however these early stages are not amenable to live imaging of haemostasis as liver development is insufficient to produce sufficient Fgb-GFP fusion protein in the *Tg(fabp10a:fgb-EGFP)*^*mi4001*^ transgenic line and there are few circulating thrombocytes visible yet in the *Tg(−6*.*0itga2b:eGFP)*^*la2*^ transgenic line (Lin, Traver et al. 2005, Vo, Swaroop et al. 2013). To facilitate live imaging with these lines we first characterised systemic infection of 5 dpf zebrafish larvae with *E. coli* strains MG1655 and UTI89. Systemic infection with 2000 CFU MG1655 did not cause appreciable mortality while a bolus of 2000 CFU UTI89 resulted in mortality starting around 1 day post infection (dpi) and continuing until 3 dpi (Figure 1A). These survival phenotypes correlated with recovery of each strain, with recoverable MG1655 burden rapidly depleting within 1 dpi (Figure 1B), while UTI89 burden peaked at 1 dpi before declining but remaining detectable until at least 3 dpi (Figure 1C).

**Figure 1:**
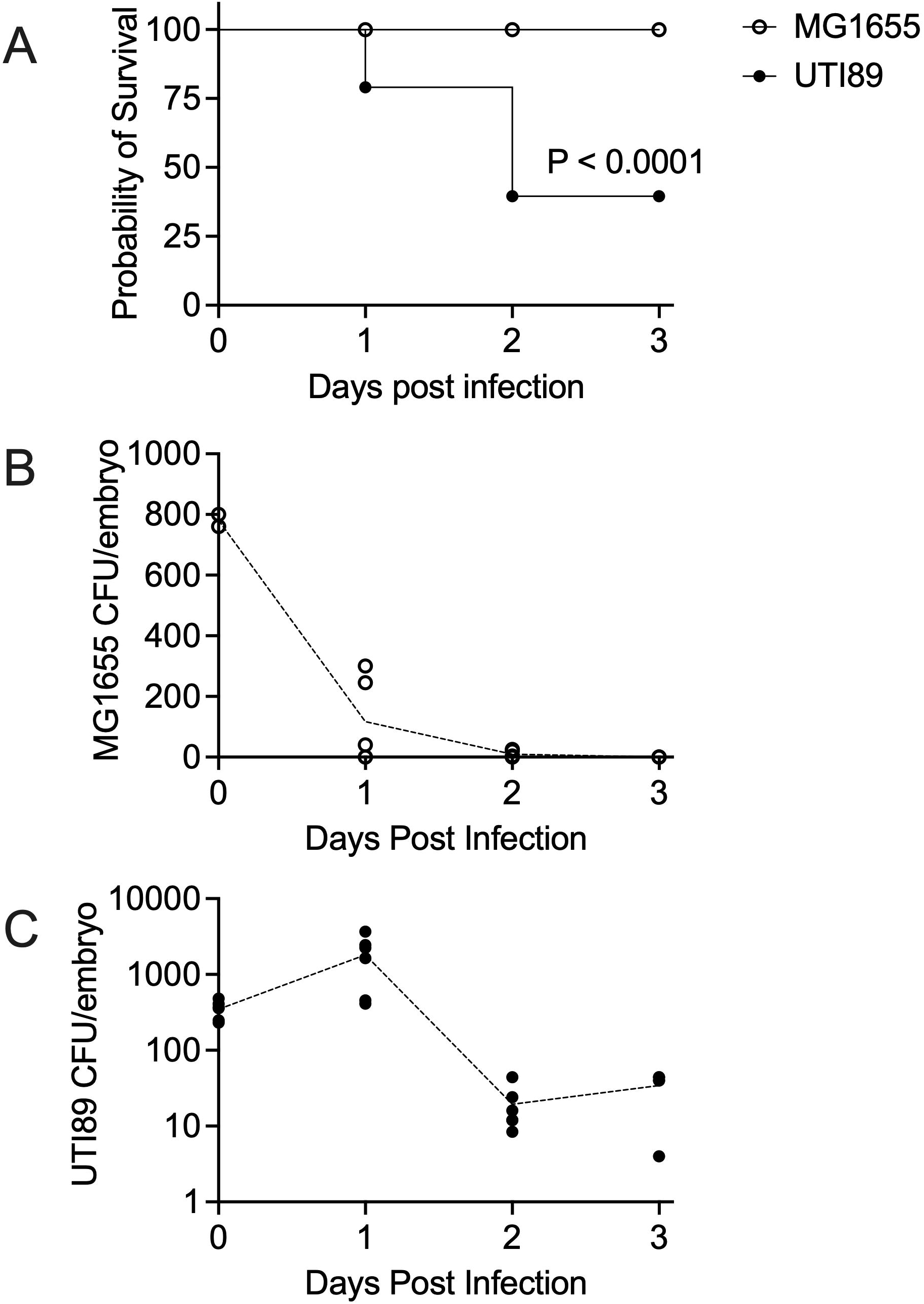
Characterisation of larval zebrafish *E. coli* systemic infection model (A) Survival analysis of 5 dpf zebrafish larvae infected with 2000 CFU MG1655 or UTI89. (B) CFU recovery from 2000 CFU MG1655-infected larvae, t0 is approximately 2 hours after infection. (C) CFU recovery from 2000 CFU UPEC UTI89-infected larvae, t0 is approximately 2 hours after infection. Data is representative of three independent experiments.

### UPEC induces coagulation in zebrafish larvae

UPEC infection-induced clotting has been observed in live rats (Melican, Boekel et al. 2008, Schulz, Chuquimia et al. 2018). To investigate if host haemostasis played a role in the superior survival of UPEC, we infected *Tg(fabp10a:fgb-EGFP)*^*mi4001*^ zebrafish larvae, where clots can be visualized by GFP fluorescence, with UTI89 UPEC carrying the pGI6 plasmid, allowing visualization of red fluorescent UPEC, and performed timelapse microscopy across the first day post infection.

We observed the progressive formation of both arterial and venous clots in infected larvae in close proximity to fluorescent UPEC (Figure 2A). We did not observe widespread clotting throughout the vasculature or clotting not associated with fluorescent UPEC. We were unable to visualise fluorescent MG1655 in the same assay, most likely due to the rapid killing of MG1655 by the host.

**Figure 2:**
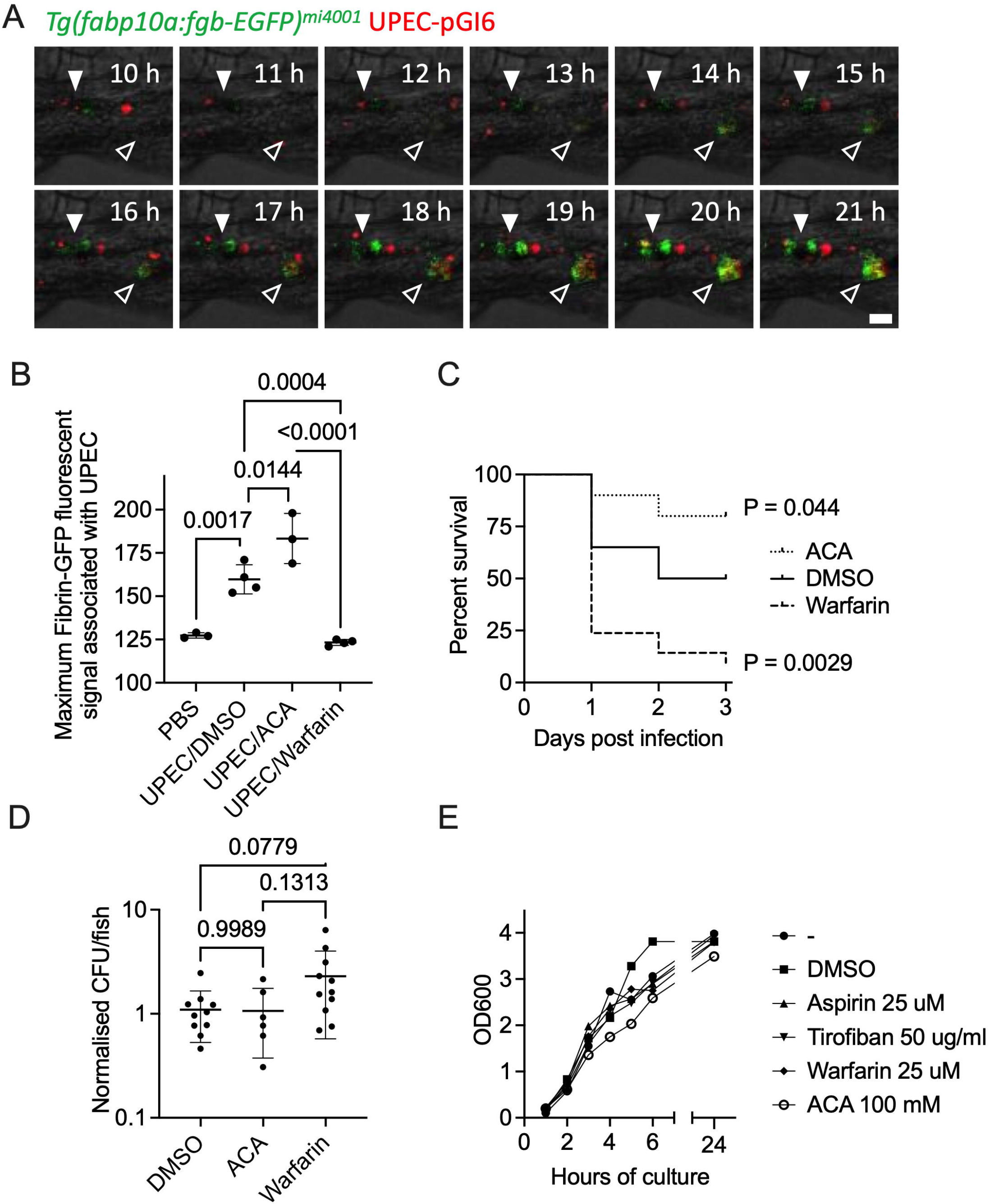
Clotting is protective against systemic UPEC infection. (A) Intravital microscopy of Fgb-GFP clot formation around UPEC UTI89 pGI6 (red) in a *Tg(fabp10a:fgb-EGFP)*^*mi4001*^ larva. Filled arrowhead indicates site of arterial clot formation, hollow arrowhead indicates site of venous clot formation. Scale bar indicates 10 μm. (B) Maximal Fgb-GFP fluorescent intensity around fluorescent UPEC from 4 hpi larvae. Each data point represents one larva. (C) Survival of 2000 CFU UPEC UTI89-infected larvae treated with warfarin or ACA. n>=20 per group, data shown is from one experiment that is representative of two biological replicates. (D) UPEC CFU recovery from 18 hpi 2000 CFU UPEC UTI89-infected larvae, data shown is pooled from two independent experiments. (E) Optical density at 600 nm readings from *in vitro* LB broth growth of UPEC UTI89 strain treated with drugs at concentrations indicated.

### Warfarin increases the severity of systemic UPEC infection in zebrafish larvae

Inhibition of clotting with heparin has been reported to worsen UPEC infection in mice (Melican, Boekel et al. 2008, Schulz, Chuquimia et al. 2018). We sought to determine if commonly prescribed anticoagulants had a similar effect in our larval zebrafish model using warfarin, a vitamin K antagonist and a commonly prescribed anticoagulant.

We have previously demonstrated that warfarin reduced mycobacterial infection-induced coagulation in zebrafish larvae (Hortle, Johnson et al. 2019), here we demonstrate that warfarin reduced Fgb-GFP fluorescence around fluorescent UPEC while aminocaproic acid (ACA) treatment stabilized clots (Figure 2B). Treatment with warfarin decreased larval survival following systemic UPEC infection while treatment with ACA conversely increased larval survival following systemic UPEC infection (Figure 2C). The decreased survival of warfarin-treated larvae correlated with increased UPEC burden at 18 hours post infection (hpi), while ACA-treated larvae had comparable levels of UPEC to control larvae (Figure 2D). Addition of warfarin and ACA to LB broth cultures of UPEC had no effect on the *in vitro* growth (Figure 2E).

### Zebrafish thrombocytes interact with UPEC

Clotting and thrombosis are coordinated during infection-induced haemostasis (Frick, Bjorck et al. 2007, Hortle, Johnson et al. 2019). To determine if zebrafish thrombocytes could play a role in controlling systemic UPEC infection, we infected *Tg(−6*.*0itga2b:eGFP)*^*la2*^ larvae, where thrombocytes can be visualized by GFP expression, with UPEC-pGI6. We observed transient interactions between zebrafish thrombocytes and clumps of UPEC, however these events were comparatively rare occurring in only 3/10 larvae imaged (Figure 4A).

### Aspirin and ticagrelor increase the severity of systemic UPEC infection in zebrafish larvae

Having shown a negative effect of inhibiting clotting, we next investigated the effect of commonly prescribed anti-platelet medications on systemic UPEC infection.

Inhibition of thrombocytes with either aspirin or ticagrelor reduced survival of infected larvae compared to DMSO-treated control larvae (Figure 4B and 4C). Treatment with either aspirin or ticagrelor increased UPEC burden at 6 hpi compared to DMSO-treated control larvae (Figure 4D). Addition of aspirin or ticagrelor to LB broth cultures of UPEC had no effect on *in vitro* growth (Figure 3E).

**Figure 3:**
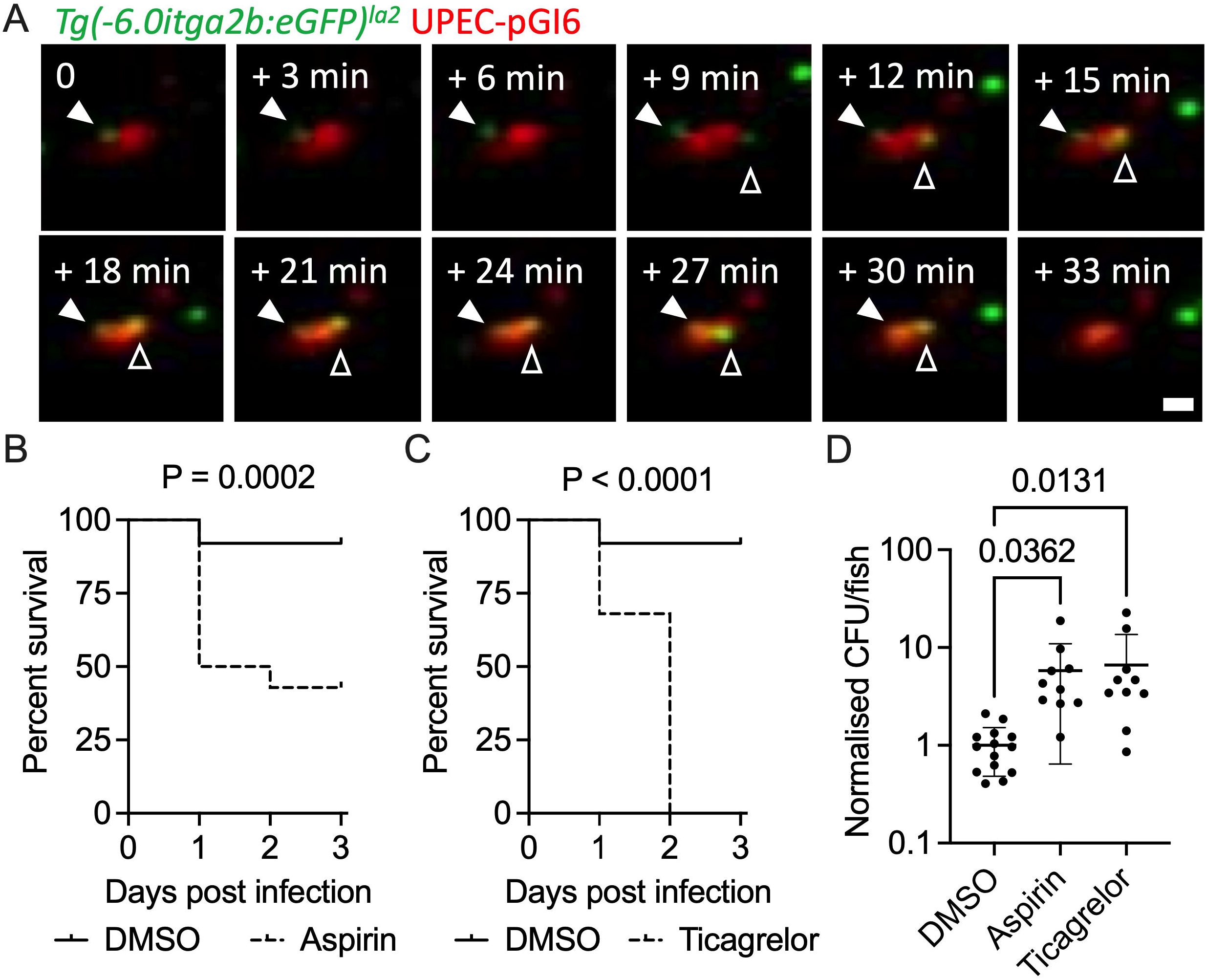
Thrombocyte activation is protective against systemic UPEC infection. (A) Intravital microscopy of green thrombocytes interacting with UPEC pGI6 (red) in a *Tg(−6*.*0itga2b:eGFP)*^*la2*^ larva. Filled arrowhead indicates first thrombocyte that interacts with clump of UPEC for 30 minutes, hollow arrowhead indicates a second thrombocyte that interacts with the same clump of UPEC for 21 minutes. Unindicated green cells are thrombocytes in circulation. t0 is approximately 1 hour post infection. Scale bar indicates 10 μm. (B) Survival of 1000 CFU UPEC UTI89-infected larvae treated with aspirin. n>=25 per group, data shown is from one experiment that is representative of two biological replicates. (C) Survival of 1000 CFU UPEC UTI89-infected larvae treated with ticagrelor. n=25 per group, data shown is from one experiment that is representative of two biological replicates. (D) CFU recovery from 6 hpi 1000 CFU UPEC UTI89-infected larvae, data shown is pooled from two independent experiments.

## Discussion

We have found that common anti-haemostatic medications worsen the survival of UPEC-infected zebrafish. Anti-haemostatic drug use is highly prevalent in the elderly as a preventative measure against heart attack and stroke. Alongside this, the risk of urinary tract infections also increases from middle to old age and there is elevated risk of recurrent urinary tract infections in aged care settings (Bennett, Johnson et al. 2016). Our data illustrates an association between preventative anti-haemostatic medication usage and the severity of UPEC sepsis.

Our results add to a growing body of literature that the haemostatic system, and specifically clotting, is crucial to the early containment of blood born UPEC (Melican, Boekel et al. 2008, Schulz, Chuquimia et al. 2018). Clinical trials of anticoagulants to treat sepsis-induced coagulopathy have delivered inconsistent results, however an emerging theme is that early coagulation is host protective while late coagulation drives pathology (Umemura, Yamakawa et al. 2016). Subsequently, administration of anticoagulants at early stage can be detrimental to the clearance of pathogen by the native immune system (Scarlatescu, Tomescu et al. 2017). There is little evidence of a beneficial effect of anticoagulants across the total range of patients with sepsis, however anticoagulants are beneficial for the treatment of critically ill subgroups with sepsis-induced disseminated intravascular coagulation (Iba, Saitoh et al. 2014, Umemura, Yamakawa et al. 2016).

Pathogen-specific effects may drive the heterogeneous host response to anticoagulant therapy. For example, Factor XI aided the containment of *Klebsiella pneumoniae* and *Streptococcus pneumoniae* in mice, while knockout of Factor XII protected mice against *K. pneumoniae* but not *S. pneumoniae* (Stroo, Zeerleder et al. 2017). The genetic conservation of vertebrate coagulation factor function and the amenability of zebrafish to rapid CRISPR-Cas9 mutagenesis suggest zebrafish could be a cost-effective platform to further delineate exact components of the coagulation cascade which we have bluntly targeted with warfarin in this study.

The negative effects of the antiplatelet drugs aspirin and ticagrelor demonstrate a host-protective role of zebrafish thrombocytes during systemic UPEC infection. Degranulation of activated mammalian platelets releases important inflammatory mediators such as antimicrobial proteins, cytokines, and ADP/ATP which can directly kill pathogens and activate cellular immunity (Deppermann and Kubes 2016). Clinical trials with antiplatelets, especially aspirin, have demonstrated an improvement in mortality of severe septic patients which, similar to anti-coagulant therapy, may reflect an effect of timing (Davis, Miller-Dorey et al. 2016). A limitation of our study is that we were unable to directly quantify thrombocyte-UPEC interactions and determine if the increased susceptibility of aspirin and ticagrelor-treated larvae was due to localized or systemic changes to thrombocyte biology.

Our CFU recovery experiments confirm warfarin treatment results in increased UPEC growth in systemic infection of zebrafish larvae. This was expected from the literature where heparin treatment facilitates increased UPEC growth in mammals (Melican, Boekel et al. 2008, Schulz, Chuquimia et al. 2018). In parallel, we observed increased UPEC burden in larvae treated with anti-platelet medications. This suggests the haemostatic system either directly controls the growth of UPEC in zebrafish or assists the zebrafish innate immune system in efforts to control systemic UPEC infection.

An important limitation of our study is that we have not established equivalency of systemic UPEC infection in zebrafish larvae with sepsis seen in mammals. Additionally, our method of infusing UPEC directly into the bloodstream of zebrafish larvae removes key steps of the natural ascending infection route via bladder colonization and pyelonephritis which account for the majority of UPEC morbidity. Zebrafish larvae are also highly tolerant of coagulopathy, as seen in an antithrombin III-deficient mutant (Liu, Kretz et al. 2014), and this may cause unexplored differences in their use of the haemostatic system to control systemic infections.

The internal concentrations of drugs achieved by immersion exposure of zebrafish larvae to drugs needs to be determined for individual substances, however previous reports have reported a range of 1-20% peak absorption of small molecules (Zhang, Qin et al. 2015, Kirla, Groh et al. 2018). Our 10 µg/ml dose of aspirin delivered by immersion exposure is likely to be at the low end of the range of peak human therapeutic plasma concentrations 2-20 µg/ml (Rosenkranz and Frolich 1985, Benedek, Joshi et al. 1995), while our 20 µg/ml dose of ticagrelor is likely to be on the high end of the peak human ticagrelor plasma concentration of 3.6 µg/ml for a 400 mg/day dose (Dobesh and Oestreich 2014), and our 33 µM, roughly equivalent to 10 µg/ml, dose of warfarin is likely to be close to therapeutic human plasma levels <1 µg/ml (Routledge, Chapman et al. 1979, Sun, Wang et al. 2006).

Future studies should investigate the interaction between the haemostatic system and the natural course of UPEC infection in both animal models and by retrospective clinical chart review. The potential for an association between aspirin and warfarin, two of the most commonly prescribed anti-haemostatic medications, with catherization-associated UPEC infections involves a significant proportion of aged care patients and may represent an important cause of excess morbidity.

## Conclusions

We find an association between the use of anti-haemostatic medication and the severity of UPEC sepsis in a zebrafish infection model.

## Funding

This work was supported by the University of Sydney Fellowship [grant number G197581]; NSW Ministry of Health under the NSW Health Early-Mid Career Fellowships Scheme [grant number H18/31086]; Centenary Institute Booster Grant to E.H.; and the Kenyon Family Inflammation Award 2017 to S.H.O., Kenyon Family Inflammation Award 2019 to E.H..

The authors have no conflicts of interest to declare.

## Acknowledgements

We thank Associate Professor Iain Duggin and Dr Bill Soderstrom (University of Technology Sydney) for the *E. coli* strains and plasmid; Dr Angela Kurz and Sydney Cytometry for assistance with imaging equipment; and Dr Vivien Chen for helpful discussion of results.

## Author contributions

E.H., and S.H.O designed the experiments. V.T., E.H., and S.H.O performed the experiments. V.T., E.H., and S.H.O wrote the paper. E.H., W.J.B., and S.H.O. supervised the project.

## Declaration of Interests

The authors declare no competing interests.

## Notes

Financial declaration: This work was supported by the University of Sydney Fellowship G197581; NSW Ministry of Health under the NSW Health Early-Mid Career Fellowships Scheme H18/31086; and the Kenyon Family Inflammation Award 2017 (S.H.O.), Kenyon Family Inflammation Award 2019 (E.H.).

### Competing Interest Statement

The authors have declared no competing interest.

### Summary of Updates

New data: CFU recovery assays, imaging Fgb-GFP +/− drug treatment Refinement of text and addition of references MG1655 comparisons in Figure 1, in vitro growth curves in Figure 2E

